# Spatial structure of synchronized inhibition in the olfactory bulb

**DOI:** 10.1101/123695

**Authors:** Hannah A. Arnson, Ben W. Strowbridge

## Abstract

Olfactory sensory input is detected by receptor neurons in the nose which then send information to the olfactory bulb, the first brain region for processing olfactory information. Within the olfactory bulb, many local circuit interneurons, including axonless granule cells, function to facilitate fine odor discrimination. How interneurons interact with principal cells to affect bulbar processing is not known though the mechanism is likely to be different than in sensory cortical regions since the olfactory bulb lacks an obvious topographical organization; neighboring glomerular columns, representing inputs from different receptor neuron subtypes, typically have different odor tuning. Determining the spatial scale over which interneurons such as granule cells can affect principal cells is a critical step towards understanding how the olfactory bulb operates. We addressed this question by assaying inhibitory synchrony using intracellular recordings from pairs of principal cells with different inter-somatic spacing. We find that in acute rat olfactory bulb slices, inhibitory synchrony is evident in the spontaneous synaptic input in mitral cells separated up to 300 *μ*m. At all inter-somatic spacing assayed, inhibitory synchrony was dependent on fast Na^+^ channels, suggesting that action potentials in granule cells function to coordinate GABA release at relatively distant dendrodendritic synapses formed throughout the the dendritic arbor. Our results suggest that individual granule cells are able to influence relatively large groups of mitral and tufted cells belonging to clusters of at least 15 glomerular modules, providing a potential mechanism to integrate signals reflecting a wide variety of odorants.

## Introduction

Inhibitory local circuits play a central role in processing olfactory information. In insects, blockade of inhibitory function in the antennal lobe, the first processing region of olfactory information, selectively impairs the normal ability of the these animals to make fine distinctions between related similar odors while leaving intact the ability to distinguish between unrelated olfactory stimuli (1). Recent work in the olfactory bulb, the mammalian equivalent to the antennal lobe, has demonstrated parallel findings when selectively perturbing inhibition onto the principal neurons mitral (MC) and tufted cells (TC) (2). These studies suggest that olfactory information can be processed through at least two distinct streams – a hardwired pathway that does not require extensive local inhibitory interactions but which reveals only relatively coarse distinctions among odors and a more complex circuit involving functions mediated by inhibitory interneurons that facilitates fine distinctions. The latter pathway may also be a key site of olfactory learning within the bulb as previous work has found both LTP and spike timing dependent plasticity on excitatory synapses onto granule cells, the primary type of GABAergic interneuron the OB (3, 4). Granule cells also receive a large proportion of top-down and neuromodulatory input (5, 6), suggesting a role of behavioral state in regulating inhibition. However, how inhibitory interneurons such as GCs function to enhance olfactory performance is not known.

Recent experimental and computational studies of OB circuitry has led to two divergent views of GC function: that GCs function through locally-mediated inhibition with minimal integration of information across processing streams, or that GCs operate by interconnecting principal cells belonging to different sensory channels, enabling the circuit to represent more complex or abstract features than either mitral or tufted cells. In the former model, GC function would likely enhance fine odor discrimination via actions on nearby principal cells belonging the same or close glomerular modules while the latter models allows for extensive cross-channel synaptic interactions. Supporting the first hypotheses are findings that local Na^+^ and Ca^2+^ spike propagation can be restricted to small subregions of the granule cell dendritic arbor (7, 8, 9, 10), physiological and computational studies that emphasize the ability of granule cells to release their neurotransmitter in a Na^+^ spike independent manner (11, 12), a physiological investigation *in vivo* demonstrating dendritic branch-specific odor tuning (13), and that GC-dependent emergent network properties such as gamma-band local field potential oscillations appeared to be uncorrelated with spiking activity in GCs (14). Alternatively, previous studies involving simultaneous intracellular recordings from pairs of MCs separated by up to 200 *μ*m, likely spanning multiple glomeruli (typically glomeruli range from 50-120 *μ*m in diameter in rodents; 15), have shown synchronous, GC-mediated inhibitory input following olfactory nerve stimulation (16) and MC depolarization by serotonin (17). It is unknown to what degree this occurs during spontaneous activity.

Through detection of TTX-sensitive coincident inhibitory postsynaptic currents (IPSCs) on principal cell pairs separated by 200 *μ*m or more, we provide evidence suggesting that dendrodendritic inhibition spans multiple glomerular columns through a mechanism that requires Na^+^-dependent action potentials. Through pharmacological manipulations, we found that the propensity for detecting coincident inhibition in MC/MC paired recordings can be increased by passive depolarization of bulbar neurons and decreased by blocking GC spiking with TTX. Phenylephrine (PE), an *α* agonist known to increase GC activity (18, 19, 20, 21), also downregulated the amount of coincident inhibition. As activation of the *α*-1R is involved in olfactory learning and memory, our finding suggests that the spatial extent of inhibitory synchrony may be differentially regulated in different olfactory behaviors and brain states. This study shows for the first time that inhibition can link a wide spatial range of glomerular columns in the absence of external synchronizing input, potentially providing a mechanism of how population codes for different odorants could be refined by local inhibitory circuits.

## Methods

### Slice Preparation

Horizontal olfactory bulb slices 300 *μ*m thick were made from ketamine-anesthetized P14-25 Sprague-Dawley rats of both sexes as previously described (22, 23). Slices were incubated for 30 min at 30°C and then at room temperature until use. All experiments were carried out in accordance with the guidelines approved by the Case Western Reserve University Animal Care and Use Committee.

### Electrophysiology

Slices were placed in a recording chamber and superfused with oxygenated artificial cerebrospinal fluid (ACSF) at a rate of 1.5 ml/min. Recordings were made between 29-32°C. ACSF consisted of (in mM): 124 NaCl, 3 KCl, 1.23 NaH_2_PO_4_, 1.2 MgSO_4_, 26 NaHCO_3_, 10 dextrose, 2.5 CaCl_2_, equilibrated with 95% O_2_/ 5% CO_2_. The K^+^ concentration was elevated in “high K ASCF” by increasing the KCl concentration to 6 mM. All whole-cell patch-clamp recordings were made with Axopatch 1C or 1D amplifiers (Axon Instruments) using borosilicate glass pipettes (WPI) of impedances ranging from 2-5MΩ pulled on a P-97 pipette puller (Sutter Instruments). Recordings were low-pass filtered at 5 kHz (FLA-01, Cygus Technology) and digitized at 10 kHz using an ITC-18 computer interface (Instrutech) connected to a PC operating Windows 7 using custom software.

Slices were imaged using IR-DIC optics on Zeiss Axioskop FS1 or Olympus BX51WI up-right microscopes. Live 2-photon imaging was performed using a custom-build laser scanning system, as described in previous publications (22, 24, 25). Neuronal cell type was determined based on IR-DIC morphology and soma laminar location. Cell type classification was confirmed in a subset of experiments using 2-photon reconstructions as described in text. Somatic separation was measured using a calibrated eyepiece reticule.

Voltage clamp recordings were made with an internal solution containing (in mM): 140 CsCl, 4 NaCl, 10 HEPES, 2 EGTA, 4 MgATP, 0.3 Na_3_GTP, 10 phosphocreatine, 5 QX-314 with a pH of 7.3 and an osmolarity of 290 mmol/kg. The cesium-chloride solution was used to reverse the chloride gradient. Alexa 594 (10 *μ*M; Invitrogen) was added to the internal solution in experiments using live 2-photon visualization. All drugs were purchased from Sigma except TTX (Calibochem) and Gabazine (Ascent). All drugs were prepared from aliquots stored at -20°C except for TTX, which was prepared from a stock solution kept at 4°C. The drugs were added to the bath by changing the external solution source.

### Data Analysis

Spontaneous IPSCs were detected and measured automatically using a custom algorithm employed in previous studies (26, 27, 28, 29). Detected events were confirmed by visual analysis. Event detection and data analysis routines were implemented in Python (version 3.5). Multiple trials (typically 10-20 episodes each lasting 10 sec; mean 16.9 trials) were acquired for each condition in each experiment. Summary data are presented as mean ± S.E.M. except where noted. Membrane variance was computed over 250 ms duration windows.

Cross-correlograms (examples shown in Figs. 2D and 4D) were computed from the onset lags between all detected IPSC pairs and analyzed between -10 to 10 ms using 0.4 ms time bins. To determine statistical significance, the maximal cross correlation value (averaged over 3 bins, 1.2 ms) reported in the actual data was compared with surrogate data created by permuting inter-IPSC intervals. The mean of 100 interval permutation runs are presented in example cross correlation plots (e.g., the grey plot in the left panel in Fig. 2D). The probability that the maximal cross correlation metric in the actual data was larger than expected by chance was computed empirically using three methods. In the first approach, we generated a distribution of 2000 maximal cross correlation values from surrogate data in which the inter-IPSC intervals were permuted. For example, if the peak cross correlation value obtained in one experiment was larger than 1900 out of 2000 cross correlation values obtained in the interval permutation runs, then probability of inhibitory coincidence would be assigned as 0.05 (100/2000). This analysis procedure was repeated five times for each experiment with the median recorded as the *interval permutation-based P value*. Throughout the study, we express randomization-derived P values as -1 * ln(P) where more positive numbers reflect lower P values. The upper limit on this metric (7.6 = -1 * ln(1/2000)) reflects experiments in which the peak cross correlation values were lower in all randomization runs than in the actual (non-permuted) data set.

**Figure 1:**
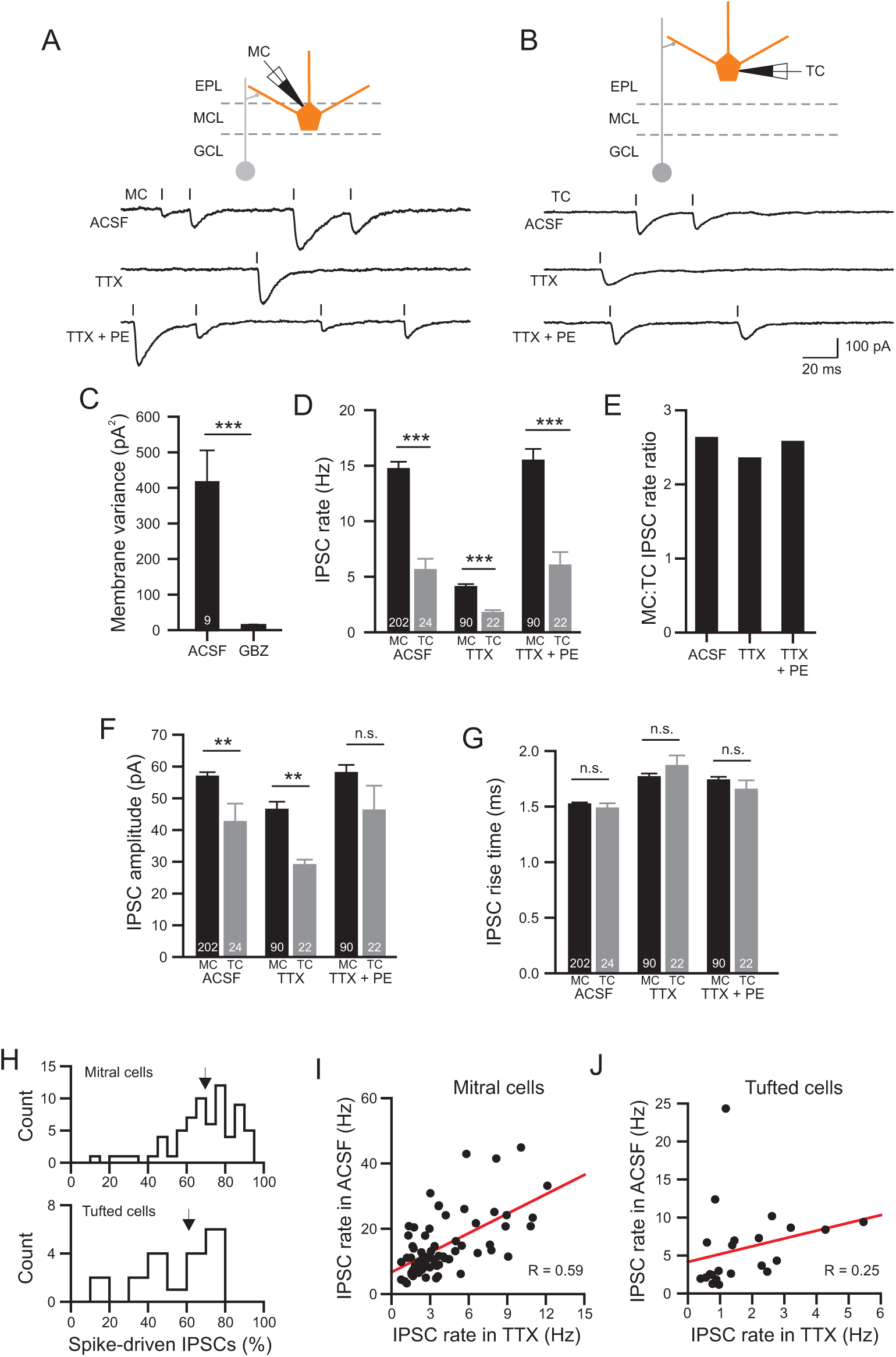
Inhibitory input to mitral and tufted cells. A, Example records of spontaneous IPSCs in ACSF (top), TTX (1 *μ*M; middle) and TTX + PE (phenylephrine; 10 *μ*M; bottom trace). Cartoon of recording configuration above traces. Vertical lines indicate automatically-detected IPSC onset times. B, Example recording illustrating spontaneous IPSCs recorded in a tufted cell under the same conditions as A. C, Plot of reduction of membrane current noise (variance) by 10 *μ*M gabazine in 9 experiments (N=5 MCs and 4 TCs). *** P = 0.003; paired t-test; t(8) = 4.2. Reduction in gabazine also statistically significant when MCs and TCs were analyzed separately (both P < 0.03). D, Plot of spontaneous IPSC rate in MCs (black bars) and TCs (grey bars) in ACSF, TTX, and TTX + PE. TC/MC comparisons *** P = 7.6 × 10^−6^ (ACSF; unpaired t-test; t(224) = 4.58), P = 1.99 × 20^−4^ (TTX; t(110) = 3.85), P = 6.11 × 10^−5^ (TTX + PE; t(110) = 4.17). Within each cell type, IPSC rates were lower in TTX than ACSF (both P < 0.002; paired t-test) but not different between ACSF and TTX + PE conditions (MCs: P = 0.54, TCs: P = 0.80). E, Plot of ratio of IPSC rates in MCs and TCs under the same three conditions. F, Plot of mean IPSC amplitude in MCs and TCs. ** P = 0.0022 (ACSF; unpaired t-test; t(224) = 3.10), P = 0.0015 (TTX; t(110) = 3.26), TTX + PE: N.S. P = 0.073. Mean IPSC amplitude also was different between ACSF and TTX conditions in MCs (unpaired t-test; P = 1.8 × 10^−4^; t(290) = 3.8) and TCs (P = 0.043; t(44) = 2.1)) but not between ACSF and TTX + PE conditions (MCs: P = 0.67; TCs P = 0.71). G, Plot of mean IPSC rise times in the same conditions (P > 0.05 in all comparisons between conditions and between MCs and TCs in the same condition). H, Histogram of percentage of TTX-sensitive (spike-driven) IPSCs compared to the IPSC rate in ACSF in 68 MCs and 22 TCs. Arrows indicate population means (MCs: 69.6%; TCs: 61.1%). I, Plot of the relation between miniature (TTX-resistant) and spontaneous IPSC rates in 68 MCs (Pearson correlation R = 0.59; P = 1×10^−7^). J, Similar plot for 24 TCs (R = 0.25; P = 0.25). In all subsequent figures * P < 0.05; ** P < 0.01; *** P < 0.005.

**Figure 2:**
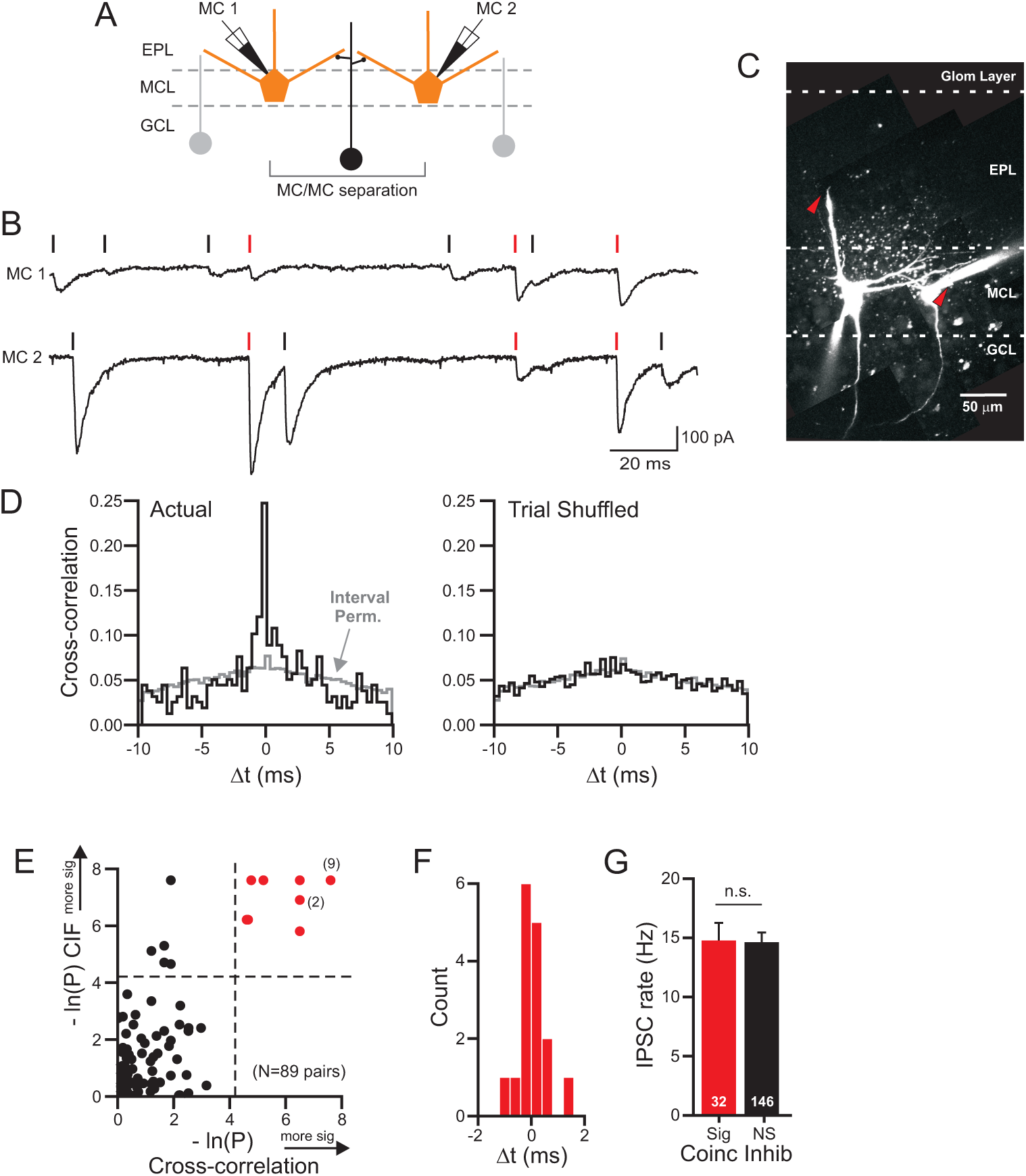
Coincident inhibitory input onto mitral cells. A, Diagram of dual MC recording configuration. B, Example simultaneous intracellular recording from two MCs. Near-coincident IPSPs (onset lags: -0.1, -0.1 and -0.1 ms) marked by red vertical lines; other automatically detected IPSCs indicated by black lines. C, 2-photon reconstruction of the two MCs shown in B. Both MCs had apical dendrites that were truncated (red arrowheads) before reaching the glomerular layer. D, Plot of cross-correlation of actual (left) and trial shuffled (right; mean of 100 runs) IPSCs times in experiment shown in B. Grey traces show chance rate of near-coincident IPSCs estimated from interval permutation in both data sets (actual and trial shuffled; plot represents mean of 100 interval permutation runs). E, Plot of the relationship between degree of inhibitory synchrony between MC/MC pairs estimated from the cross correlation method shown in D (the *dual randomization P value* described in the Methods) and the clipped cross intensity function (CIF). Probability of near-coincident inhibition increases with higher numbers (axes reflect -1 * natural log(P value)). Red symbols reflect P < 0.015 (4.2 = -1 * ln(0.015)). Multiple experiments with similar P values indicated by “(n)”. F, Histogram of IPSC lag corresponding to maximal cross correlation in 16 MC/MC paired recordings with statistically significant inhibitory synchrony (0.4 ms bins). G, Comparison of mean IPSC rates in MC/MC pairs with (red) and without (black) inhibitory synchrony (P = 0.94; unpaired t-test).

**Figure 3:**
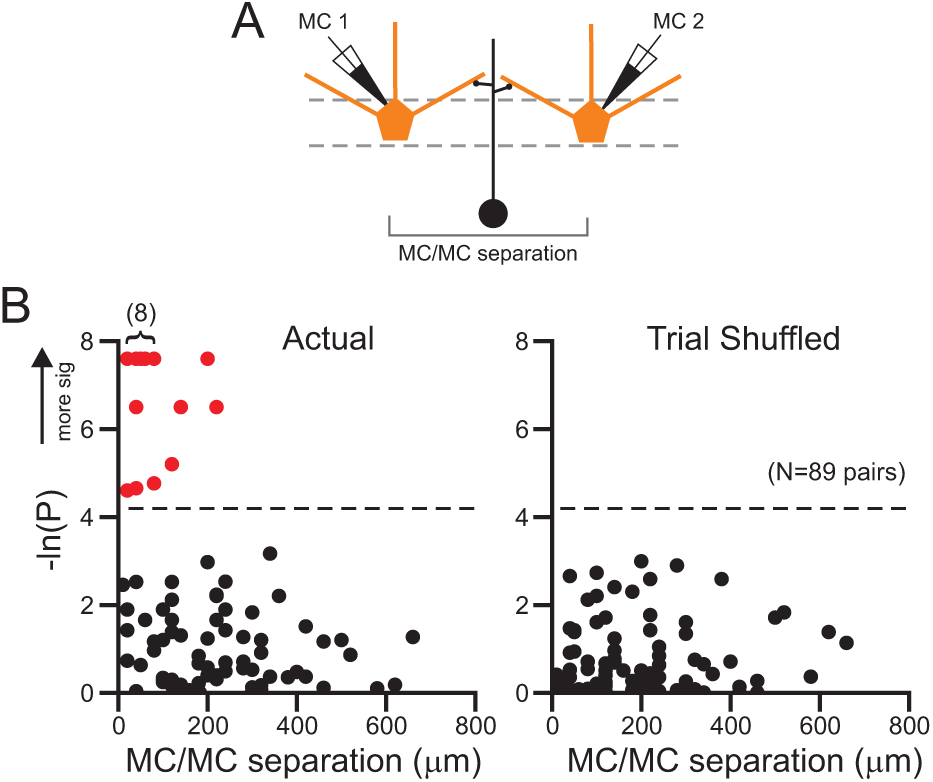
Spatial scale of coincident inhibition among mitral cells. A, Diagram of dual MC recording configuration. B, Plot of the probability of inhibitory coincidence versus separation between MC somata in 89 experiments from actual IPSC timing (left) and following trial shuffling (right). Red symbols indicate P < 0.015 (> 4.2 on -1 * ln(P) axis; dashed line). Cluster of red dots (marked “(8)” in left plot corresponds to 8 experiments with similar P values.

**Figure 4:**
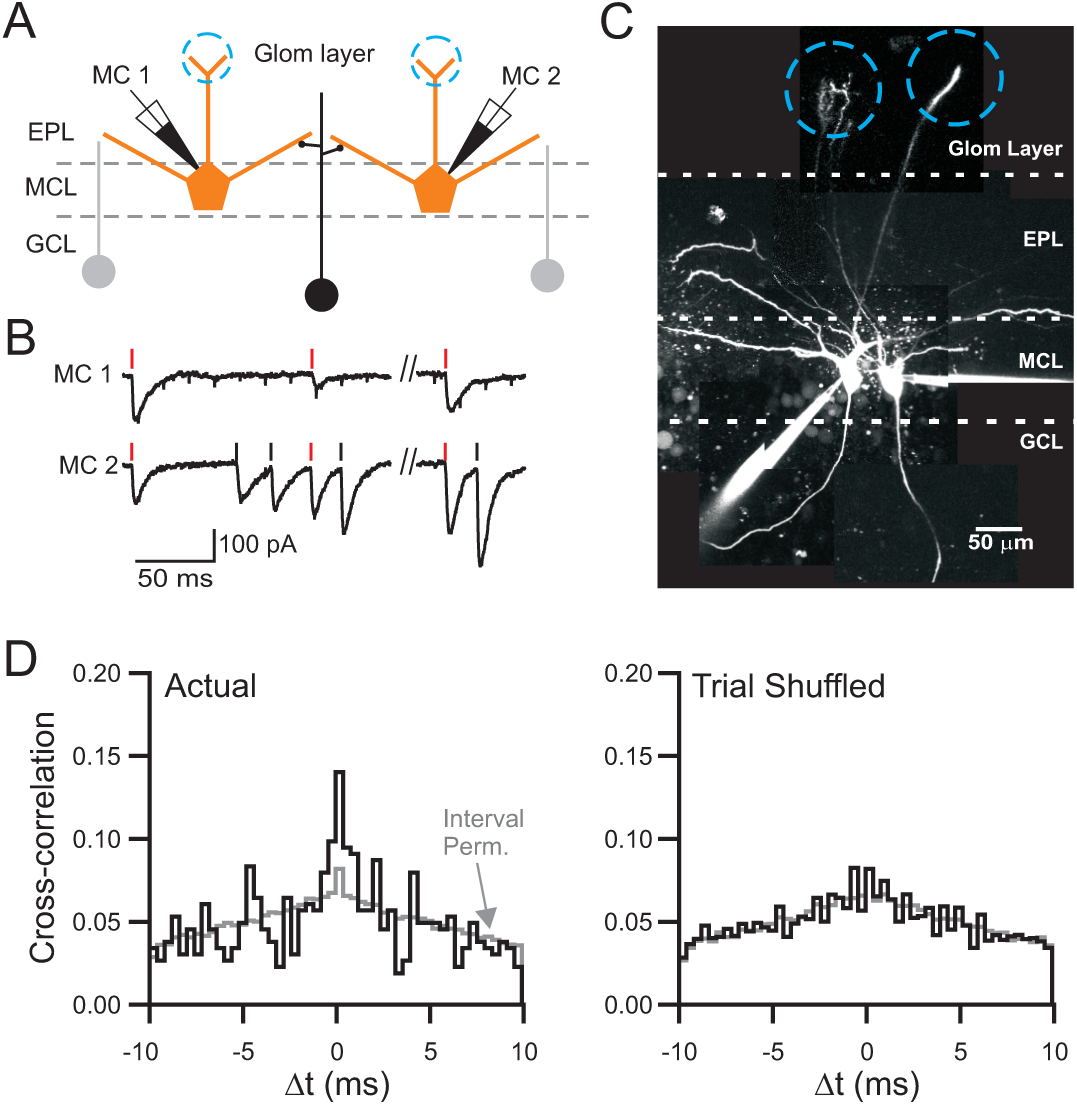
Coincident inhibition in mitral cells belong to different glomeruli. A, Diagram of recording configuration. B, Example recording from two MCs with near-coincident IPSCs (onset lags: -0.1, -0.6 and -0.2 ms). C, 2-photon reconstruction of the two MCs shown in B. The apical dendrites terminated in different glomeruli separated by 130 *μ*m (indicated by blue circles). D, Plot of cross correlation in the MC/MC pair shown in B-C based on actual (left) and trial shuffled (right; mean of 100 trial shuffling runs) IPSC onset times. Grey traces show the shift predictor estimated from interval permutation (100 interval permutation runs).

In the second randomization control method, we used trial shuffling to estimate whether each experiment had beyond-chance rates of inhibitory coincidence. In this test, we randomly selected different trials for cell A and for cell B from the same experiment, creating at data set composed exclusively of non-simultaneous records (eg, trial 1 from cell A compared with trial 3 from cell B). This process was repeated 2000 times in each experiment to generate a P value based on the distribution of trial shuffled recordings. As with the interval permutation control procedure, we repeated the trial shuffling analysis five times and recorded the median of those runs as the *trial shuffling-based P value*. Both the interval permutation and trial shuffling control procedures abolished the peak near 0 ms present in experiments with obvious inhibitory synchrony determined by visual inspection of raw intra-cellular records. Figures 2 and 4 illustrate cross correlation plots following interval permutation and trial shuffling on two example paired recordings with frequent near-coincident IPSCs.

Both interval permutation and trial shuffling approaches generated similar metrics expressing the degree of inhibitory synchrony evident our primary data set of 89 paired MC/MC recordings assayed under standard ACSF conditions (R = 0.92; P < 10^−30^; correlation performed on the -ln(P) metric as used throughout the study). We also computed a *dual randomization P value* where we combined both randomization methods by computing bootstrap distributions based on interval permutations of trial shuffled data. Using a standard 0.05 threshold for statistical significance, the *dual randomization P value* generated the most conservative ranking of inhibitory synchrony across experiments (N = 17 experiments with empirical P values lower than this threshold), compared with N = 21 for the interval permutation approach and N = 23 using only trial shuffling. 16 experiments had P values < 0.05 in all three randomization tests. Rather than report separate results from interval permutation and trial shuffling procedures, we only report only the final *dual randomization P value* throughout study and in the illustrations.

We set a threshold for statistical significance at 0.015 using the *dual randomization P value* throughout the study. This value balanced competing goals of minimizing the false rejection rate while controlling the family-wise error rate. A more conservative approach, such as Bonferroni corrections, would adjust the significance threshold to 0.05/N where N is the number of independent tests performed. This simple approach proved unworkable since the very low thresholds in our large N data set (P = 0.00056 for our population of 89 MC/MC paired recordings) rejected many experiments with obvious inhibitory coincidence and central peaks on cross correlation plots that were abolished by both interval permutation and trial shuffling. As noted in other work, this approach often rejects true positives and, therefore, tends to reduce statistical power (30). Employing a Bonferroni correction approach also would lead to different thresholds for statistical significance across the different groups of experiments. The empirical P value approach we followed has been used in previous studies (31, 32, 33, 34) and allowed us to specify a uniform threshold (0.015) that was more stringent than the P value traditionally employed with single tests (0.05). Using all three randomization approaches, we found a large drop off in P values near this threshold (from < 0.0101 in all experiments labeled significant to 0.042 in the best experiment labeled as non-significant). In our primary data set with 89 paired MC/MC recordings, using this threshold resulted in 16 experiments with significant inhibitory synchrony. In Fig. 2E, we also estimated the degree of inhibitory synchrony using the clipped cross-intensity function approach (CIF; 35) we have used in a related study (17). This metric was based on the count of near-coincident IPSP (onset times within 3 ms) divided by the total number of IPSCs in the cell with the fewest IPSC. As with the cross-correlation method, we computed P values for each experiment empirically based on a distribution of CIF metrics generated in surrogate data in which inter-IPSC intervals were permuted.

The estimates of the rates of above-chance inhibitory coincidence presented in Fig. 7D were computed by subtracting the rates of near-coincident IPSCs (onset latencies within 3 ms) before and after trial shuffling. We plotted the rate of near-coincident IPSCs beyond expected by chance as a percentage of the TTX-sensitive IPSC rate in each cell. For example, if the actual rate of near-coincident IPSCs was 2 Hz which then dropped to 0.5 Hz following trial shuffling, then the rate of “beyond-chance” near-coincident IPSCs would be 1.5 Hz. If the spike dependent rate of spontaneous IPSCs was 15 Hz in one of the cells in the paired recording (reflecting 20 Hz spontaneous IPSCs recorded in ACSF and 5 Hz in TTX), then “beyond chance” coincident IPSCs would constitute 10% of the spike-driven spontaneous IPSCs in that neuron.

**Figure 5:**
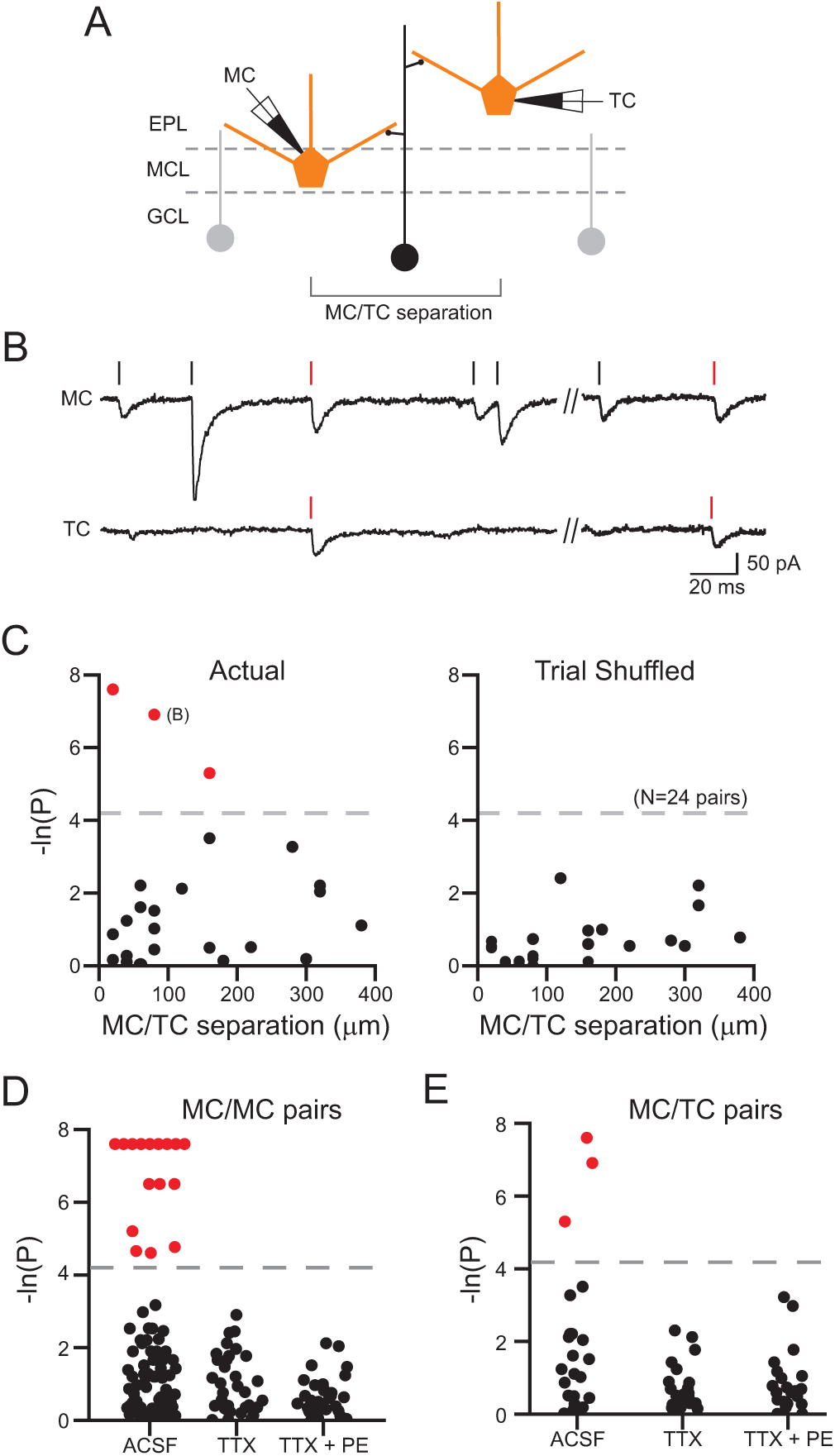
Coincident inhibition in mitral/tufted cell paired recordings. A, Diagram of recording configuration. B, Simultaneous MC (top trace) and TC (bottom) recording showing near-coincident IPSCs (onset lags: -0.1 and -0.9 ms). C, Plot of the relationship between the degree of inhibitory synchrony and somatic separation in 24 MC/TC paired recordings based on the actual (left) and trial shuffled (right) IPSC onset times. Three experiments with statistically significant inhibitory synchrony (P < 0.015; dashed lines) denoted by red symbols. X axis reflects only the component of the somatic separation along the MCL. D, Plot of the degree of inhibitory synchrony in MC/MC paired recordings assayed in ACSF (N=89), TTX (N = 43) and TTX + PE (N=43). Experiments with statistically significant inhibitory synchrony (P < 0.015; dashed line) denoted by red symbols. E, Similar plot for MC/TC paired recordings (N=24 experiments in ACSF, N=22 in TTX and TTX + PE).

**Figure 6:**
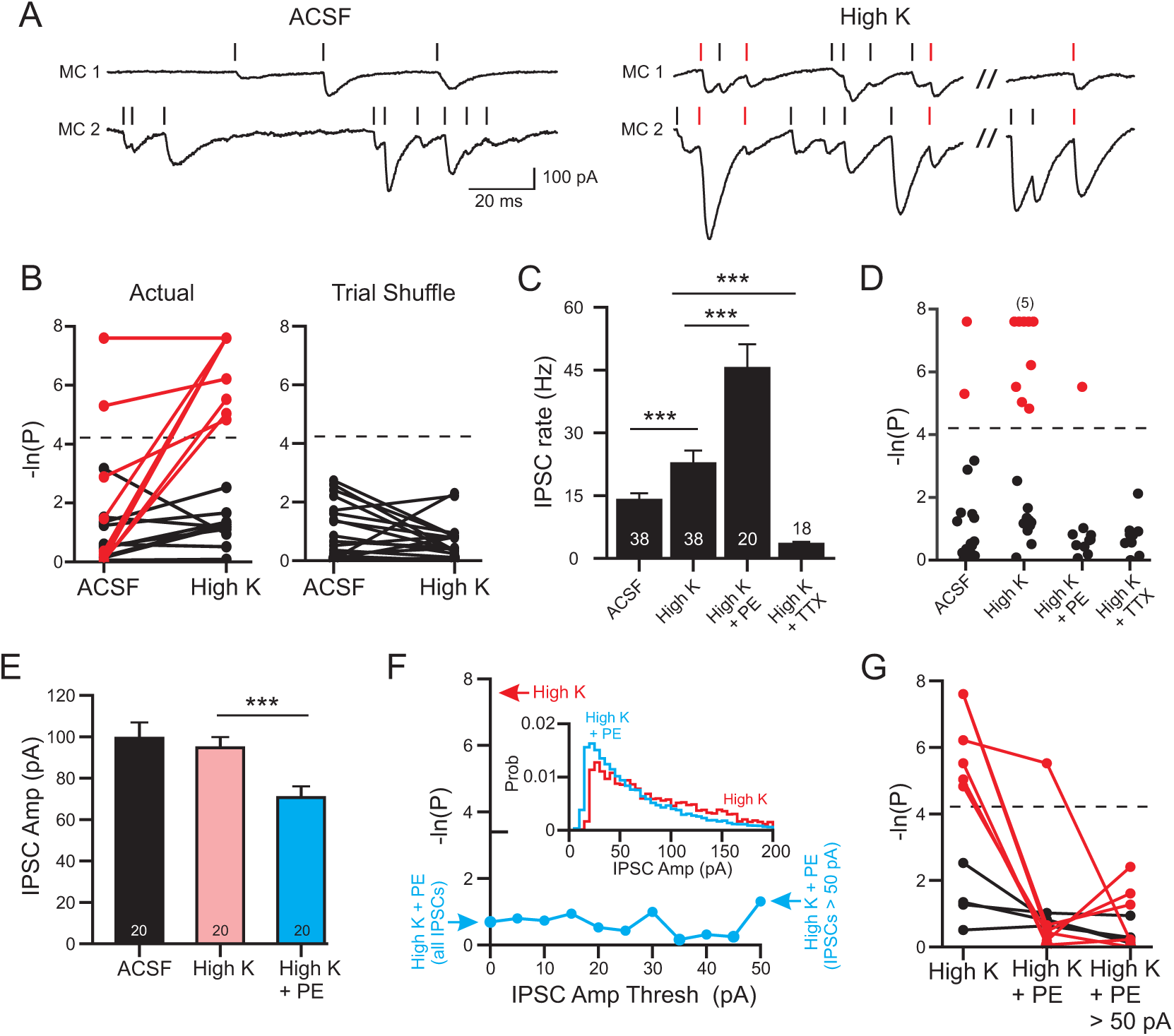
Passive depolarization with elevated K^+^ increases inhibitory synchrony. A, Example dual MC recording with near coincident IPSCs (onset lags: -0.4, -0.4, -0.3 and 0.1 ms) in high K ACSF but not in control ACSF. B, Plot of degree of inhibitory synchrony in 19 paired MC recordings tested in control and high K ACSF (P values based on actual IPSC onset times shown in left panel and following trial shuffling on right). Experiments with statistically significant inhibitory synchrony in high K ACSF indicated by red lines (7 experiments with P < 0.015 only in high K ACSF and 2 experiments with P < 0.015 in both conditions). C, Plot of mean IPSC rate in following passive depolarization using high K ACSF. *** (ACSF vs. high K ACSF: P = 0.0001, t(37)= 4.35, paired t-test; high K vs. high K + PE: P = 0.00044, t(56)=3.74; unpaired t-test; high K to high K + TTX: P = 0.00011, t(54)=4.17, unpaired t-test). D, Plot of the degree of inhibitory synchrony in MC/MC paired recordings following passive depolarization with high K ACSF. Dashed line represents same P = 0.015 threshold for statistical significance used throughout study. Cluster of 5 experiments with similar P values indicated by “(5)”. E, Plot of mean IPSC amplitude following passive depolarization. *** P = 0.0035 (t(19) = 3.11, paired t-test) F, Plot of degree of inhibitory synchrony versus the lower cutoff of IPSC amplitudes analyzed in one MC/MC paired recording in high K + PE. Far right point represents results of cross correlation analysis including only IPSCs > 50 pA in both MCs while far left point reflects the cross correlation analysis when including all detected IPSCs. This paired recording showed statistically significant inhibitory synchrony in high K ACSF (red arrow by top of Y axis) Inset, histogram of IPSC amplitudes in this example paired recording in high K (red; N = 5419 IPSCs) and high K + PE (blue; N = 18588 IPSCs) conditions. G, Plot of the degree of inhibitory coincidence following passive depolarization with high K ACSF, Middle column represents experiments in high K + PE when all IPSCs are included while right column represents high K + PE when only IPSCs > 50 pA are included. Dashed line at P = 0.015.

**Figure 7:**
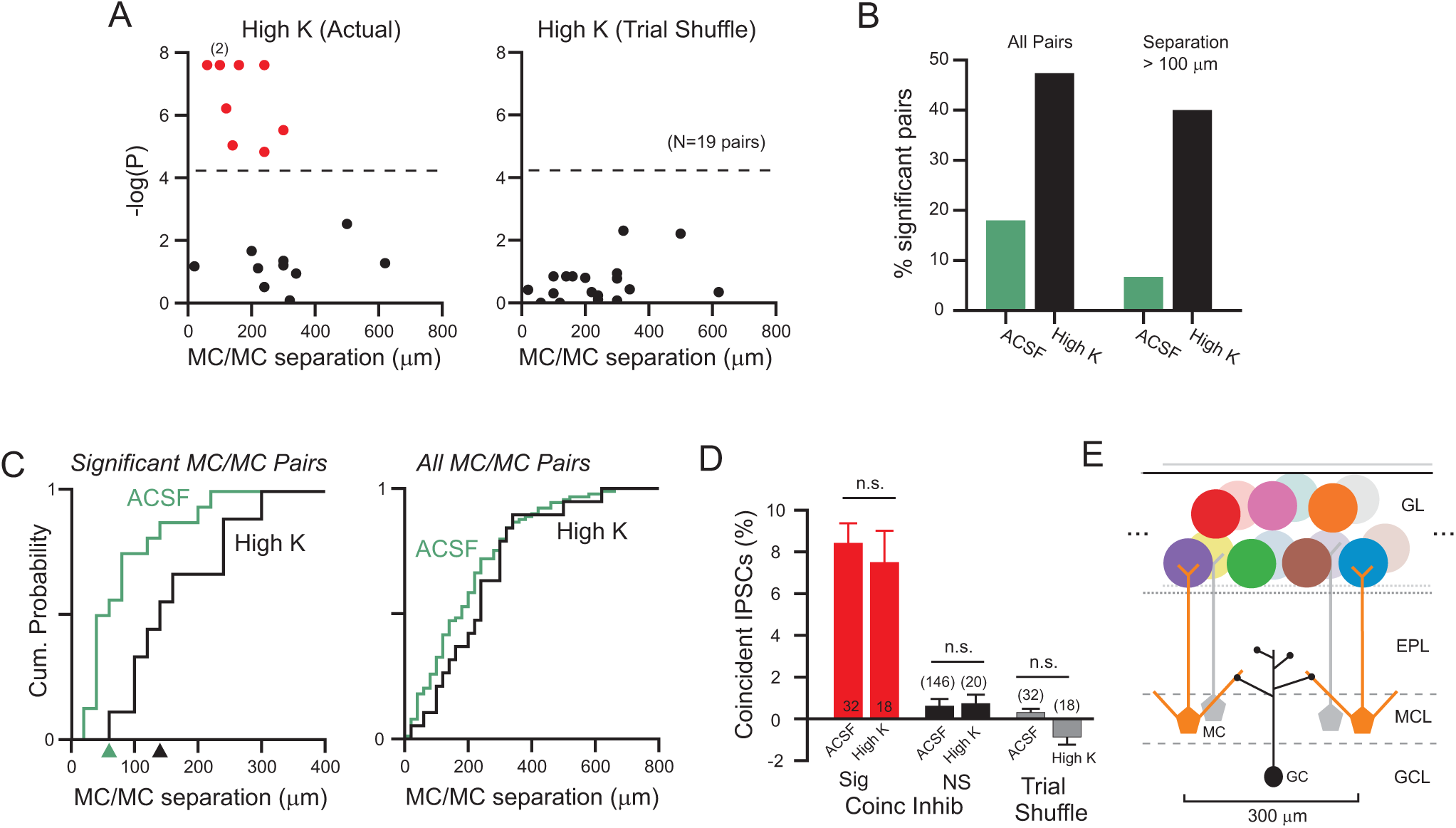
Passive depolarization expands spatial scale of coincident inhibition. A, Plot of the relationship between degree of inhibitory coincidence and somatic separation in 19 MC/MC paired recordings based on actual (left) and trial shuffled (right) IPSC onset times. Two experiments with similar P values indicated by “(2)” in left plot. Dashed line at P = 0.015. B, Comparison of the fraction of MC/MC paired recording experiments with statistically significant inhibitory synchrony (P < 0.015) in control (green bars) and high K ACSF (black). Left bars represent all MC/MC paired recording experiments while right pair of bars represents only experiments where MC cell bodies were separated by > 100 *μ*m. C, Plot of cumulative distribution of MC somatic separations in control (green) and high K ACSF (black). Plots on left restricted to MC/MC paired recordings with statistically significant inhibitory synchrony (N = 16 ACSF and 9 high K; curves different, P = 0.0486, D(19,19)=0.421, K-S test) while plot on right reflects all MC/MC experiments (N = 89 ACSF and 19 high K). Arrowheads by X axis in left plot indicate MC somatic separation cutoffs that include half of the relevant population (analogous to EC50 estimates). D, Plot of percent of all IPSCs that reflect “excess” coincident IPSCs (beyond the fraction of near-coincident IPSCs (onset lag < 3 ms) expected by chance) in experiments with statistically significant inhibitory synchrony (red bars), experiments without demonstrable inhibitory synchrony (black bars) and in all MC/MC paired recordings following trial shuffling (grey bars). All comparisons between bar pairs did not differ significantly (all P > 0.05, unpaired t-test). Cartoon of expected fan-out from individual bulbar interneurons (example GC illustrated). See *Discussion* for details.

### Cellular reconstructions

The horizontal span of the GC dendritic arbor was calculated from 19 published GC images from the following references: 36, Figure 2, 37 Figures 1C, 2A, 2H and 3A, 8 Figure 2, 38 Fig 1, 39 Fig 4A, 5A and 6A, and 40 Figure 1M-O. To account for shrinkage during fixation, the measured width was increased by 15% in papers that explicitly stated that fixation was used (41 – 10% lateral shrinkage, 42 – 10-20% on each dimension). The mean GC dendritic width we estimated from these published studies was 113 ± 11.1 *μ*m with a standard deviation of 48.5 *μ*m.

We estimated the thickness of the glomerular layer in 10 OB slices visualized using a 5X objective (185 ± 21 *μ*m; mean ± S.D.). We estimated the glomerular packing density using our measured GL thickness and the mean glomerular diameter reported in two anatomical studies (75 *μ*m; 43, 44). The horizontal span of MC lateral dendrites was estimated from maximal 2-photon Z-stack projections (N=33 lateral dendrites from 21 visualized MCs; 349 ± 180 *μ*m; mean ± S.D̤. The actual length of the lateral dendrites was ∼10% larger than the horizontal span (the component of distance within the MCL), reflecting angled trajectory of these dendrites from the cell body into the EPL.

## Results

We divided our analysis of inhibitory input onto principal cells in this study into two sections, starting with determining the properties of spontaneous ISPCs in MCs and TCs (Fig. 1) and then focusing on measuring coincident inhibition across pairs of intracelluarly-recorded principal cells to assess the spatial scale of inhibitory synchrony in the OB (Figs. 2-7). Both MCs and TCs receive frequent spontaneous IPSCs, shown as inward currents in the example voltage-clamp recordings in Fig. 1A-B which employed CsCl-based intracellular solutions that reversed the normal chloride gradient. Spontaneous inwards currents were abolished by the GABA-A receptor antagonist gabazine (10 *μ*M; 96.6% reduction in membrane current variance; Fig. 1C), indicating that spontaneous IPSCs form the majority of the detectable synaptic input onto OB principal cells under these recording conditions. Most of the detected IPSCs in MCs appeared to originate from synapses onto secondary dendrites and not from synapses onto the distal apical dendrite. In a subset of 21 MCs visualized using live 2-photon microscopy, we found only a small decrease in the rate of spontaneous IPSCs in MCs with apical dendrites truncated before they reached the glomerular layer (12.7 ± 1.5 Hz; N=15) compared with MCs with intact apical dendrites (14.6 ± 1.6 Hz; N = 6: P > 0.05; unpaired t-test; mean ± S.E.M.).

We first compared the baseline frequency and amplitude of inhibitory inputs onto both classes of principal cells. Mitral cells received a significantly higher rate of spontaneous IPSCs than TCs (Figure 1D-E), consistent with results from a recent comparison of bulbar neurons (45). The mean IPSC amplitude was modestly smaller in TCs than MCs (Fig. 1F) while the rising phase kinetics were similar in both cell types (Fig. 1G). In both cell types, blockade of voltage-gated Na^+^ channels with 1 *μ*M TTX dramatically decreased spontaneous ISPC frequency (middle panel in Fig. 1D), suggesting that most synaptic responses resulted from Na^+^–based spiking in GCs or other classes of GABAergic interneurons. Enhancing GC excitability by stimulating *α* adrenegic receptors with phenylephrine (PE; 10 *μ*M; 18, 19, 20, 21) was able to restore basal IPSC rates in both principal cell types even in the presence of TTX (right panel in Fig. 1D).

Surprisingly, throughout these manipulations the ratio of spontaneous IPSC frequencies between MCs and TCs remained relatively constant (Fig. 1E), suggesting that the two cell types differed primarily in the density of inhibitory connections with GCs. The proportion of spike-driven IPSCs also was similar in both cell types (mean of 70% in MCs vs 61% in TCs; Fig. 1H). We observed a strong correlation between miniature IPSC rates and basal spontaneous IPSC rates that was more pronounced in MCs (Fig. 1I; r = 0.59; P < 10^−7^) than TCs (Fig. 1J). As with the other measures of inhibitory function, we found more variability among TC IPSC rates, including an outlier recording with very high spontaneous IPSC rates. Excluding this outlier TC, the correlation between mini IPSC rates and basal spontaneous IPSC rates was statistically significant (r = 0.532, P = 0.013; r = 0.251 including outlier value). Together, these results suggest that the basal inhibitory synaptic tone in MCs and TCs is determined primarily by the density of GABAergic synapses on the secondary dendrites (and potentially somatic sites). Both cell types likely sample from similar pools of spontaneously active interneurons, accounting for the similar properties of individual IPSCs in MCs and TCs, the similar fraction of TTX-sensitive IPSCs and the ability of miniature IPSC rates to predict the basal spontaneous IPSC rates. The difference in inhibitory synaptic tone we and others (45) observe in OB principal cells likely reflects fewer synaptic contacts on TCs vs MCs, consistent with previous anatomical studies (46; 47).

### Coincident inhibition between pairs of OB principal cells

After determining baseline properties of spontaneous IPSCs in MCs and TCs, we next recorded from pairs of principal cells (N = 89 MC/MC and 24 MC/TC) separated by variable distances to determine the spatial scale of over which synchronized inhibition occurs in the OB. A subset (∼15%) of paired MC recording had multiple coincident IPSCs evident by visual inspection of intracellular records, such as the example paired recording in Fig. 2A-C. We used two independent control procedures to determine if the rate of coincident IPSCs was greater than expected by chance in each paired MC/MC recording. We first computed the cross-correlation based on IPSC onset times in both cells and compared the maximal cross-correlation function (“actual”) to randomized data in which the intervals between recorded IPSCs were permuted (“interval permutation”; Fig. 2D, left). In the second analysis method, we compared the maximal cross-correlation function computed from actual data with the same analysis performed trial shuffled recorded (Fig. 2D, right; see Methods), generating a dataset with physiological IPSC timing statistics but no actual coincidences since only non-simultaneously recorded datasets were analyzed (e.g., trial 1 from MC-A with trial 2 from MC-B). We computed the probability that actual maximal cross correlation function was larger than expected by chance by combining the interval permutation and trial shuffling procedures 2000 times; the threshold for significance employed was P < 0.015; see Methods for details). 16 of 89 MC/MC paired recordings met this criterion for above-chance rates of near coincident IPSCs. All 16 MC/MC paired recordings identified as having inhibitory synchronization assayed through the cross-correlation method also exhibited statistically significant rates of coincident IPSCs when tested using the cross-intensity function method employed in previous work (Fig. 2E; 26).

Near-coincident synaptic inputs identified by our analysis (which captured onset latencies that differed by up to 20 ms) were nearly synchronous in the subgroup of 16 MC/MC paired recording classified as statistically significant. The mean latency between IPSC onset times across the two cells was 0.05 ± 0.13 ms (range -1.0 to 1.4 ms; N=16 experiments; histogram shown in Fig. 2F). There was no difference between the overall mean IPSC rates in the 16 significant MC/MC paired recordings and the other 73 paired recordings (Fig. 2G). The inhibitory synchrony we observe was unlikely to arise from glomerular-layer local circuits since the 16 significant MC/MC paired recordings included 3 experiments in which both MCs were reconstructed using live 2-photon imaging and at least one MC had a truncated apical dendrite that was severed within the EPL (including the example MC pair shown in Fig. 2A-C).

We determined the spatial scale of coincident inhibition by plotting the P value computed from the interval permutation analysis against the separation between the two MC somata (Fig. 3A-B). Most MC/MC paired recordings with significant inhibitory synchrony occurred between closely-spaced neurons (inter-soma distances <100 *μ*m). The maximal somatic separation that generated inhibitory synchrony beyond chance levels in standard ACSF was 220 *μ*m. When restricted to inter-somatic spacing less than 220 *μ*m, we still observed no difference in the mean IPSC rate between significant and non-significant MC/MCs pairs (P=0.71; unpaired t-test; t(57) = 0.38) suggesting that our metric of inhibitory synchrony was not simply revealing an underlying bias toward MCs with high (or low) spontaneous IPSC rates. The distribution of P values estimated from interval permutation in trial shuffled data (all less than 0.015; Fig. 3B, right; equivalently, greater than 4.2 on the scale in Fig. 3B) was similar to the distribution of the actual non-significant MC/MCs (non-trial shuffled data; black symbols in Fig. 3B, left). These results suggest our dataset of MC/MCs paired recordings comprised two populations: a minority (18%) which received coincident inhibition, presumably reflecting divergent output from one or more spontaneously spiking interneurons, and MC/MC pairs that received no detectable inhibitory coincidence.

The relatively large spatial scale over which we observe inhibitory synchrony under basal conditions (up to 220 *μ*m somatic separation) suggests that individual interneurons can couple MCs associated with different glomeruli (mean glomerular diameter 75 *μ*m; 43, 44). We confirmed this was possible by recording and visualizing two MCs with intact apical dendritic tufts that terminated in nearby but different glomeruli (Fig. 4A-C; inter-soma distance 40 *μ*m, apical dendritic tufts separated by 130 *μ*m). The cross correlation function in this paired recording (Fig. 4D, left; P < 0.0005) had a peak at 0.2 ms.

Under basal (unstimulated) conditions, we also observed inhibitory synchrony between heterogeneous pairs of principal cells (experiments including one MC and one TC recording; Fig. 5A). Figure 5B illustrates an example of coincident IPSCs recorded in a MC/TC pair with inter-soma spacing of 80 *μ*m (measure as horizontal distance through the MCL, ignoring the different distances in the MCL-to-GL direction). Using the same criteria applied to MC/MC paired recordings, 3 of 24 MC/TCs had statistically significant inhibitory coincidence (Fig. 5C), including the example recording presented in Fig. 5B. The three significant MC/TC pairs were separated by between 20 and 150 *μ*m (span along the MCL). The mean latency between near-coincident IPSCs was 0.2 ± 0.33 ms (range -0.2 to 1 ms). As with the MC/MC paired recording that showed elevated rates of near-coincident IPSCs, there was no difference in the mean IPSC rates between MC/TC pairs with and without demonstrable inhibitory synchrony (P > 0.05; unpaired t-test; t(46) = 1.73). These results suggest that not all inhibitory local circuits in the OB are segregated into MC- and TC-specific subgroups. Instead, individual non-glomerular layer interneurons appear to innervate both populations of principal cells.

### Spike-dependent inhibitory synchrony

We next asked if near coincident IPSCs recorded in pairs of principal cells required Na^+^-based action potentials in the common presynaptic interneurons. Bath application of TTX (1 *μ*M) abolished inhibitory synchrony in all 16 MC/MC pairs with statistically significant rates of near coincident IPSCs (Fig. 5D). No previously non-significant paired recording experiments became significant through this treatment. The loss of inhibitory synchrony did not simply reflect the lower overall rate of spontaneous IPSCs in TTX since addition of the GC-stimulating *α* receptor agonist PE (10 *μ*M; 18, 19, 20, 21), failed to promote synchronous inhibition in MC paired recordings (Fig. 4D; 0/44 experiments). We found similar results in heterogeneous MC/TC pairs where TTX abolished inhibitory synchrony in the 3 paired MC/TC recordings with statistically significant rates of near coincident IPSCs (Fig. 4E). Since voltage-gated Na^+^ channels were already blocked in MC and TC voltage-clamp recordings (because of QX-314 in the internal solution), these results suggest that synchronous IPSCs resulted from spontaneous spiking in GABAergic interneurons that form divergent synaptic connections onto MC and MC/TC ensembles.

If near coincident IPSCs recorded in pairs of principal cells reflect spontaneous spiking in presynaptic GABAegic interneurons, we hypothesized that passively depolarizing interneurons by elevating extracellular K^+^ concentration (from 3 to 6 mM; “high K” ACSF) would enhance inhibitory synchrony. As shown in the example records in Fig. 6A, high K ACSF increased both the rate of spontaneous IPSCs in MCs and also the incidence of near-coincident IPSCs. Of the 19 paired MC experiments in high K ACSF, 9 (47%) had statistically significant inhibitory synchrony, including the paired recording shown in Fig. 6A. Two of the 9 significant MC/MC recordings already had significant rates of inhibitory synchrony in control conditions, prior to treatment with high K ACSF (Fig. 6B). In the remaining 7 significant MC/MC recordings, no inhibitory synchrony was evident except in high K ACSF. The mean IPSC latency in the 9 MC paired recordings with near coincident IPSCs in high K ACSF was 0.24 ± 0.19 ms (range -0.6 to 1.4 ms).

Our results thus far demonstrate divergent effects of passive depolarization with high K ACSF, which promoted inhibitory synchrony, and activation of adrenergic receptors with PE, which stimulated GABA release (increasing miniature IPSC rates) but did not trigger more frequent coincident IPSCs. Consistent with this model, PE combined with high K ACSF increased spontaneous IPSC rates (Fig. 6C) but abolished most inhibitory synchrony (only 1/10 experiments with PE + high K ACSF has statistically significant inhibitory synchrony; Fig. 6D). As with our earlier results suggesting a requirement for Na^+^-based action potentials, inhibitory synchrony present in high K ACSF was abolished in TTX (Fig. 6D; 0/9 experiments including 3 MC paired recordings that had statistically significant inhibitory synchrony in high K ACSF).

While high K ACSF did not affect the amplitudes of spontaneous IPSCs recorded in MCs, PE triggered a modest but statistically significant decrease in mean IPSC amplitude (29% decrease; Fig. 6E). This difference primarily reflected an over-representation of IPSCs with very small amplitudes (see IPSC amplitude histograms in Fig. 6F, inset). We, therefore, tested whether the lack of statistically significant inhibitory synchrony following treatment with PE and high K ACSF reflected a “dilution” effect that prevented detecting above-chance inhibitory synchrony. We tested for this possibility by restricting our cross correlation analysis to spontaneous IPSCs greater than an arbitrary minimum amplitude. As shown in plot in Fig. 6F, we failed to find statistically significant inhibitory synchrony when analyzing different subsets of large-amplitude IPSCs in the 10 MC pairs with no evident inhibitory synchrony in high K ACSF. Eliminating spontaneous IPSCs less than 50 pA also abolished inhibitory synchrony in the one MC/MC paired recording with statistically significant rates of near-coincident IPSCs (Fig. 6G), suggesting that some synchronous inhibitory inputs generated relatively small amplitude (< 50 pA) IPSCs.

Passively depolarizing bulbar neurons with high K ACSF also increased the spatial extent of inhibitory synchrony. As shown in Fig. 7A, MC pairs with inter-soma distances up to 300 *μ*m exhibited statistically significant inhibitory synchrony (versus up to 220 *μ*m in control ACSF). As in previous control analyses, trial shuffling eliminated all statistically significant inhibitory synchrony (Fig. 7A, right). The overall proportion of MC/MC paired recordings with significant inhibitory synchrony increased from 18% in control ACSF to 47% in high K ACSF (Fig. 7B). We found an even more pronounced increase in the proportion of experiments with significant inhibitory synchrony when only considering MC pairs with inter-soma separations >100 *μ*m (right bars in Fig. 7B). Consistent with this hypothesis, we found a statistically significant difference between plots of cumulative inter-soma distance in experiments with inhibitory synchrony in control and high K ACSF (Fig. 7C, left). The midpoint in MC separations with inhibitory synchrony (similar to EC50 estimates) shifted from 60 *μ*m in control to 140 *μ*m in high K ACSF (arrowheads along the X axis in Fig. 7C, left) There was no difference in the cumulative inter-soma distance plots when considering all MC/MC pairs (regardless of whether they had inhibitory synchrony) between control and high K ACSF (P > 0.05; K-S test; K-S (19,19) = 0.15; Fig. 7C, right).

While high K ACSF increased the fraction of experiments with inhibitory synchrony and the spatial extent of inhibitory synchrony, this treatment did not affect the proportion of putative synchronized IPSCs within each recording. Actual measurements of near-coincident IPSCs (e.g., number of IPSCs in one recording with a corresponding IPSC within 3 ms in the other, simultaneous recording) reflect the sum of two different frequencies: the rate of real (“biological”) coincident IPSCs, presumably corresponding to coordinated release of GABA at two different synapses from the same interneuron, and the rate of chance coincidences. We calculated the “excess coincidence rate”–reflecting an estimate of the underlying frequency of actual coordinated GABA release events–by subtracting the rate of random coincidences (computed by permuting inter-IPSC intervals) from the measured rate of near coincident IPSCs. The excess inhibitory coincidence rate modestly increased from 0.70 ± 0.14 Hz in the 16 MC/MC with statistically significant inhibitory synchrony in control ACSF to 1.62 ± 0.69 Hz in the 9 MC/MC pairs with inhibitory synchrony recorded in high K ACSF (P > 0.05; unpaired t-test). However, expressed as a proportion of the ongoing spontaneous IPSC frequency in each condition, there was almost no difference in the rate of excess IPSCs between control (8.4% of all spontaneous IPSCs) and high K ACSF (7.5%; Fig. 7D). As expected, we found near-zero rates of excess coincident IPSCs following trial shuffling or when assaying MC/MC pairs that did not have statistically significant inhibitory synchrony (Fig. 7D, right columns).

The similar percentage of near-coincident events detected under different levels of rates of ongoing spontaneous IPSCs suggests that inhibitory synchrony reflected inputs from small but constant fraction of presynaptic inputs. This 8 percentage estimate could arise from a single spontaneously active interneuron that innervated both MCs assuming each MC also receives input from ∼12 other active interneurons (or from 2 convergent interneurons and ∼23 other active interneurons, etc). This percentage would remain constant during passive depolarization tests if the treatment had a similar effect in all presynaptic GCs. If the inhibitory coincidence we report arises on average from a single active interneuron, then this provides an estimate of the total number of active presynaptic interneurons each MC receives (∼13).

## Discussion

In this study, we assayed functional synaptic connectivity patterns using paired intracellular recordings and make three primary conclusions. First, we find that the frequency of spontaneous IPSCs is strongly correlated with the frequency of miniature TTX-resistant IPSC in MCs suggesting that most inhibitory inputs to MCs behaves as a uniform population. Second, we find that Na^+^ spike-driven coincident inhibition can link pairs of principal cells belonging to different glomerular columns as well as different types of principal cells (mitral and tufted cells). These results provide the first direct demonstration that distant pairs of principal cells (with apical tufts innervating different glomeruli) can be functionally linked by coincident inhibition under physiological conditions. This finding is consistent with previous results from Schoppa (16) who observed inhibitory coupling between distant mitral cells following tetanic electrical stimulation. However, direct pathway stimulation can generate synchronize firing in multiple interneurons–potentially generating coincident inhibitory postsynaptic responses in MCs as a consequence. These types of “common driver” mechanisms are unlikely to account for inhibitory synchrony apparent under resting (non-stimulated) conditions employed in the present study, leaving spontaneous Na^+^ spiking in GCs that are presynaptic to both MCs as the most likely explanation for above-chance rates of coincident IPSC. And finally, we found that coincident inhibition onto MCs depends on Na^+^ spiking in GCs and can be disrupted by strong depolarizing stimuli. Together, these results suggest that bulbar interneurons can play a pivotal role in the olfactory system by coordinating responses in principal cells belonging to different glomerular channels. These synaptic interactions can function to modulate mitral and tufted cell firing patterns to reflect contextual information associated with the larger glomerular network activated by an odor.

### Difference in inhibitory tone between mitral and tufted cells

To our knowledge, our study is the first to compare inhibitory tone in mitral and tufted cells under normal physiological conditions and when Na^+^-based spiking is abolished by TTX. Our finding that TCs receive less pronounced inhibition (approximately three-fold fewer IPSCs) is consistent with several previous reports that assayed the rate of spontaneous inhibitory postsynaptic responses (45; 48). This different in spontaneous IPSC/P rate could be explained by preferential innervation of MCs by more active interneurons compared with TCs or by a difference in overall density of inhibitory innervation in the principal cell types. Our observation that the three-fold difference in IPSC rate between MCs and TCs is maintained when Na^+^ spiking is blocked with TTX and then when GC excitability is enhanced by PE suggests that this difference reflects primarily a higher density of inhibitory synaptic inputs on MCs. In principle, the difference in synaptic number may reflect the narrower span of secondary dendrites of TCs compared to MCs (47, 46). However, direct tests of postsynaptic responses elicited by focal GABA uncaging along the secondary dendrites (49) demonstrates that MCs (and presumably TCs) are primarily sensitive only to inhibitory inputs contacting the proximal section of the secondary dendritea (estimated ∼ 150 *μ*m, twice the 75 *μ*m electronic length constant for dendritic GABA-evoked responses recorded in the somata; 49). Differences in the length of the secondary dendrites between MCs and TCs beyond this proximal region are unlikely to influence the incidence of coincident inhibition detected in somatic recordings.

Secondary dendrites of MCs and TCs also arborize in different zones within the EPL where they contact different subpopulations of GCs (50, 46). Diminished inhibitory tone in TCs could, therefore, reflect a lower density of reciprocal contacts formed between these two GCs and principal cells. While Greer and colleagues (51) found a similar density of reciprocal synapses throughout the EPL, to our knowledge the density of dendrodendritic synapses formed by deep and superficial GCs has not be been compared quantitatively. Our finding that coincident inhibition can be detected in MC/TCs (in addition to MC/MC pairs) suggests that the distinction between deep and superficial GCs targeting different subclasses of principal cells is not absolute with some GCs likely forming synaptic connections between MCs and TCs (and see 50 and 52).

### Synchronized inhibition links principal cells across multiple glomerular columns

We detected synchronized inhibition using standard methods (34) based on cross-correlation analysis of IPSC onset times and employed two different control procedures to estimate the frequency of coincident IPSCs expected by chance in each experiment (interval permutation and trial shuffling). The spontaneous IPSC frequency was only modestly reduced in visualized MCs with apical dendrites truncated before reaching the glomerular layer, compared with intact MCs, suggesting that most of the inhibitory tone detected in somatic recordings from OB principal cells originates from interneurons outside of the glomerular layer, presumably GCs. While several populations of non-GC interneurons exist in the GCL, these appear to target either TCs (GL-dSACs, 53) or GCs (25, 54). A recent study (55) has found that parvalbumin (PV)-positive interneurons located in the EPL form reciprocal synapses with MCs and, therefore, could also contribute to the inhibitory synchrony we observe. However, PV EPL interneurons appear to fire at high rates (> 40 Hz in 55), which is inconsistent with the very low rates of beyond-chance synchronized IPSCs in our MC/MC paired recordings (∼0.7 Hz in ACSF).

Coincident inhibition onto MC/MC and MC/TC pairs was abolished in TTX, suggesting that they likely originate from GABA release sites coupled by Na^+^-based action potentials in GCs. The absence of detectable inhibitory synchrony in our study was unlikely to occur simply because of the lower IPSC rate in TTX since we also observed no detectable coincident inhibition (beyond chance levels) when GCs were stimulated with PE after Na^+^ spikes were abolished. This treatment restored near-control rates of spontaneous IPSCs in both MCs and TCs but did not reveal inhibitory synchrony. This finding suggests that in absence of Na^+^-based APs, direct depolarization of GCs is unable to trigger coordinated GABA release at multiple dendritic spines.

Granule cells generate APs in response to depolarizing stimuli that propagate throughout the dendritic arbor (56). Since many of MC pairs we find receive synchronized inhibition are separated by distances that are the far extreme of what should be possible given the electrotonic length constant of MC secondary dendrites assayed from focal GABA uncaging (∼75 *μ*m; 49), it is likely that Na^+^-based APs coupled GABA release at different spines along the GC dendritic tree. This mechanism would explain the ability of TTX to abolish inhibitory coupling in MC/MC and MC/TC paired recordings and the inability of increasing excitability in TTX with PE to recover inhibitory synchrony in pairs known to share a common presynaptic interneuron.

Functional (13) and computational (57) studies have suggested an important role for spatially-localized Na^+^ and Ca^2+^ spikes in GCs (8, 9, 10). The extent of spike propagation within GC dendrites represents an attractive mechanism to regulate the degree of inhibitory synchrony in principal cells. At the smallest extreme, individual GC spines are likely able to release GABA autonomously, leading to an increase in asynchronously inhibition in MCs. Removing extracellular Mg^2+^ greatly facilitates GABA release from GCs by enhancing Ca^2+^ influx through NMDA receptors located in dendritic spines (58, 26). However, this treatment does not lead to an increase in coincident IPSPs, even between nearby pairs of MCs (26)–likely reflecting uncoordinated release events in different spines. In contrast to low Mg^2+^ treatment, the present study found that increasing extracellular K^+^ increased both IPSC frequency and the incidence of coincident inhibition. Presumably elevated K^+^ increased the frequency of spontaneous spiking in GCs and, therefore, the frequency of near-simultaneous release of GABA on spines that synapse onto different principal cells. Further increasing the excitability of GCs by combining elevated K^+^ with PE greatly diminished the incidence of synchronized inhibition. This effect probably reflected depolarization blockade of AP generation in GCs rather than dilution of synchronized IPSCs by an elevated frequency of small-amplitude asynchronous events since restricting our analysis to large-amplitude IPSCs failed to recover evidence for coincident inhibition.

### Functional significance of coincident inhibition onto OB principal cells

The central finding in this study is that individual GCs can generate coordinated inhibition onto relatively distant principal cells. This result suggests that GCs can function to influence firing patterns of output neurons belonging to different glomerular columns. While we demonstrate directly that coincidence inhibition occurs spontaneously in MCs associated with nearby glomeruli (Fig. 4), the spatial extent over which we observe synchronized inhibition suggests that individual GCs influence activity over an even larger range of glomerular columns. The average diameter of a glomerulus is ∼75 *μ*m (range 50-120 *μ*m; 15) and a rough estimate of the 2-dimensional packing density is 2.5 glomeruli per 100 *μ*m span parallel to the MCL (185 *μ*m estimated glomerular layer width). Assuming at least two layers of intact glomeruli per slice (but likely 3-4 in some slices), leads to a maximal span of 14 glomerular columns per GC (Fig. 7E) and an estimated average connectivity of 7 columns per GC. This analysis is based solely on detecting the incidence of coincident IPSCs exceeding chance, so it likely represents an underestimate since infrequently discharging GCs will generate too few coincident IPSCs to be detected in our assay. This analysis also does not account for the variable trajectory of the apical dendrite as it passed through the EPL, which also could enhance the effective glomerular span of GC-mediated inhibition.

Because there is very little chemotopic structure within the glomerular layer (59, 60, 61, 62), it is possible for an individual GC to affect the firing patterns of principal cells responding to a wide range of odorants. Inhibitory postsynaptic responses can trigger rebound spikes in MCs (63, 64, 65, 16), providing a potential mechanism to generate a cohesive neural signal shaped by GC activity, consisting of coincident APs across a subpopulation of OB principal neurons that could then be detected in downstream piriform cortical neurons. Coordinated inhibition may also function to sculpt MC and TC firing patterns to enhance subtle differences evoked by sensory input, faciliating decorrelation of related odor responses (66, 67, 68, 69, 70). Since most MC/GC and TC/GC synaptic connections in the EPL are reciprocal (12, 71), our results suggest that individual GCs also receive excitatory input from a relatively large range of sensory neuron subtypes, helping to explain the complex sensory responses observed in GC recordings *in vivo* (72, 73, 67, 74, 75, 61, 38).

